# Microstructure evolution of multivalent ion-complexed polyelectrolyte gels

**DOI:** 10.1101/2025.11.10.687581

**Authors:** Yu Sen Jie, Di Jia

## Abstract

Ions perform crucial functions in biological systems. However, the understanding of the relation between polyelectrolyte gel and ion is confined to the effects of swelling by salt solutions on macro- and micro-properties of gel. Here we explore the microstructure evolution of gel with changing crosslinking density and concentration of types of salts measured by dynamic light scattering (DLS). The dissociation-association of the ion bonds will induce a stretched exponential mode in the correlation function. The stretched exponent *β* reflects the subchain dynamics of gel strands and undergoes a transition from 0.33 to 0.5 when the multivalent salt concentration increases to the vicinity of phase separation boundary. It demonstrates that the Gaussian chain-like gel strands with Rouse dynamics transform into molten-globule-like strands with Zimm dynamics at high multivalent salt concentration.

## 1 Introduction

Biomacromolecules in living organisms are closely associated with ions through dynamic bonds. In particular, the multivalent ions play their roles based on not only the net charge but also their molecular nature. These ions deeply take part in various physiological processes including metabolism, osmotic regulation, catalysis, specific recognition *etc*.^1-3^. For example, the volume explosion of mucus is driven by the exchange of Ca^2+^ and Na^+^.^4^ Zn^2+^ constitutes the catalytic or structural center of approximately 10% of human proteins.^5^ DNA and RNA require ions including Mg^2+^ to screen their charges, thus enabling nucleic acid-protein interactions.^6^ Organic multivalent ions spermine^4+^ is involved in cellular metabolism found in all eukaryotic cells.^7^

Many biological systems behave like hydrogels, such as mucus, nerve cells, and the extracellular matrix, which reconstruct their structures in response to the physical interactions (electrostatic interaction, dipolar interaction, hydrophobic and hydrophilic interactions, hydrogen bonding, and van der Waals forces) with environment, thereby performing diverse functions.^8-10^ However, biological hydrogels are featured with their complexity and difficulty to control key parameters like polymer volume fraction, charge density and crosslinking density. Accordingly, an idealized model system can offer insight into the association between the gel matrix and ions. Polyacrylate gel is a standard system due to its controllable physical parameters and especially for tuning its charge density, several methods can be employed including simply polymerizing acrylate (possibly also acrylamide or acrylic acid),^11-14^ or hydrolyzing the pendant amide groups of polyacrylamide gel.^15^ These previous researches focus on the effects from salt solutions swelling on the macro- and microscopic properties of gels. Through changing crosslinking density and swelling by monovalent salt NaCl solution, polyacrylate gels with different volume fraction *ϕ* can be obtained. The independent measurements combined with mean field theory show scaling laws among moduli *G* and gel diffusion coefficient *D*, which are *G* ≈ *ϕ*^2^, *D* ≈ *ϕ*^2/3^.^13^ For the gel swelling in divalent salt solution, a discontinuous and reversible change in volume with varying salt concentrations is observed, which is called the volume phase transition. It is reported that when the gel swells in a solution containing both Na^+^ and Ca^2+^, the concentrations of the cations vary smoothly in the gel despite the abrupt change in the gel volume. Moreover, by changing Ca^2+^ concentration, the elastic modulus *G* of gels and volume fraction *ϕ* follow a defined power law of *G*∼*ϕ*^1/3^, indicating that Ca^2+^ does not alter the effective crosslinking density.^14^ Data from small-angle neutron scattering further verify that Ca^2+^ does not affect the large-scale (low *q* range) where the chemical crosslinks primarily generate intense scattering.^16^ The volume phase transition induced by Ca^2+^ is mainly attributed to the influence on the thermodynamics of the solution which reduces the repulsion between the adjacent chains, and the resulting formation of fiber-like bundling chains.^11, 12^

Multivalent salt solution swelling of the gel systems has been studied extensively, but how gel systems organize themselves in the presence of multivalent ions has long been overlooked. Especially, polysaccharides, proteins and other biomacromolecules incorporate types of multivalent ions into their processing, modification, formation of 3D functional structure, and gelation into network.^17^ The present work provides a systematic investigation into the evolution of microstructure of gel in the presence of multivalent ions by using dynamic light scattering (DLS). Recently, it was discovered a hierarchical dynamical power law of fluctuations in gels involving physical bonds, with stretched exponent *β* = 1/(2*υ* + 2) when the subchain dynamics obeys the Rouse dynamics, and for Gaussian chain statistics with ν = 0.5, it has *β* = 1/3.^18^ While for Zimm dynamics, it has *β* = 1/(3*υ* + 1).^19^ Such a stretched exponential mode represents the dissociation-association hierarchical relaxations of the reversible physical crosslinks. In this work, we expand this universal law to polyacrylate-multivalent ion gel systems. Through polymerizing monomers and chemical crosslinker within multivalent ion solutions, the as-prepared gels exhibit stretched exponential decay in DLS with *β* equaling to 1/3 to 1/2. The exponent *β* reflects the gel strands changing conformation from flexible random-walking state at low salt concentration or high chemical crosslinking density, to compacted and collapsed state at high salt centration or low chemical crosslinking density. Here it is the first time to report the microstructure evolution of polyelectrolyte gels in the presence of multivalent ions.

## 2 Materials and methods

### 2.1 Materials

Acrylamide (Am) (40 wt%), UV initiator α-ketoglutaric acid, and spermine were bought from Sigma-Aldrich. Crosslinker bis-acrylamide solution (MBAA) (2 wt%), calcium chloride (CaCl_2_), magnesium chloride (MgCl_2_), aluminum chloride (AlCl_3_) hexahydrate, and barium chloride (BaCl_2_) were purchased from Macklin. Sodium acrylate (Ac) was purchased from Energy Chemical and was prepared as 20 wt% stock solution. Hydrogen chloride (HCl) was purchased from Sinopharm. Deuterium oxide (D_2_O) was purchased from J&K Scientific. Dimethyl sulfoxide (DMSO) was purchased from Concord Technology. Deionized water was obtained from a Milli-Q water purification system (Millipore). The resistivity of deionized water used was 18.2 MΩ·cm. Hydrophilic polyvinylidene fluoride (PVDF) filters with 220nm pore were purchased from Millex Company.

### 2.2 Preparation of poly(acrylamide-acrylate) (PAm-PAc) gel with multivalent salt

PAm-PAc gel was synthesized by free radical polymerization. The charge density ([Ac]/[Am+Ac]) was fixed at 80%, the total monomer concentration was fixed at 1 M, and the concentration of initiator was 1.5% (molar fraction of total monomers). Three parameters were tuned here, which were the chemical crosslinking density, the salt concentration, and the types of salt cations. Spermine was neutralized by HCl to pH = 6.86, to ensure it carrying four positive charges.^20^ The synthesis procedure for the PAm-PAc gel with 0.5% crosslinking density is shown as an example. 0.355 mL of Am (40 wt%), 3.762 mL of Ac (20 wt%), 0.385 mL of MBAA (2 wt%), 21.9 mg initiator and 4.998 mL deionized water were put into a tube. The pre-gel solution was bubbled with nitrogen gas for 20 min to remove any dissolved oxygen. The polymerization was irradiated with UV light for 4 hours.

### 2.3 Dynamic light scattering (DLS) measurement

The DLS tubes were washed with deionized water and acetone separately more than three time. The tubes were dried in the oven overnight, and then we used aluminum foil to wrap them. Distilled acetone through an acetone fountain setup was used to clean up these tubes. All the stock solutions were filtered through 220 nm PVDF hydrophilic filter before used. The preparation of all the samples should be conducted in a super clean bench. DLS measurement was performed on a commercial spectrometer (ALV/CGS-3), which was equipped with a multi-τ digital time correlator (ALV/LSE-5004) using a wavelength of 532 nm laser light source. For each sample, the intensity at the different scattering angles was correlated, and each data point was obtained by averaging over three samples. All the experiments were conducted at 23 °C.

DLS measures the intensity-intensity correlation function *g*_2_(*q, t*) by means of a multi-channel digital correlator and related to the electric field correlation function *g*_1_(*q, t*) through the Siegert relation:^21^

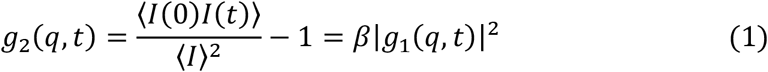

where *t* is the decay time and *I*(*t*) is the scattered intensity, *β* is the instrument coherence factor. *g*_1_(*q, t*) can be represented as a sum or integral over a distribution of relaxation rates *Γ* by

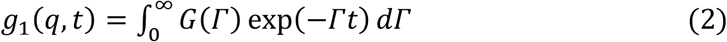

If the mode is diffusive in nature, then the decay rates are linear with *q*^2^ for all the scattering angles. From the slope of *Γ* = *Dq*^2^ (*q* = (4πn/λ) sin(θ/2) denotes the scattering wave vector with scattering angle θ, wavelength of the incident light λ and refractive index of the medium n) across all *q*, diffusion coefficient *D* is obtained. The characteristic relaxation rate *Γ* was analyzed by using CONTIN method and multiple exponential fitting method at each angle.^21, 22^ For the functions with multiple modes, the correlation function *g*_1_(*q, t*) can be fitted by a sum of exponential decays. The function *g*_1_(*q, t*) was fitted by ORIGIN software and the error between the raw data and the fitted function was minimized after iterations were performed. Residuals of the difference between the fitted curve and the raw data are randomly distributed about the mean of zero and the residuals do not have systematic fluctuations about their mean value.

### 2.4 ^1^H nuclear magnetic resonance spectrometer (^1^H NMR)

^1^H-NMR was performed by Bruker Avance Ⅲ 400 HD spectrometer. The samples of pre-gel solution and the gel were synthesized in 90% H_2_O and 10% D_2_O. The DMSO was used as external standard substitute with 0.1 mol/L. The concentration of monomers in the pre-gel solutions and the concentration of unreacted monomers remaining in the gels were determined by the integral area ratio of vinylic protons of the monomers versus DMSO protons.

### 2.5 Zeta potential measurement

Zeta potential measurement was performed by Malvern Panalytical Zetasizer Pro. All the samples were prepared without MBAA and diluted to half before put into sample cells to ensure mobility and avoid generating bubbles.

### 3 Results and discussion

For gels, the electric field correlation function *g*_1_(*q, t*) is related to the correlation of the gel strands displacement along the longitudinal direction *u*_*l*_, expressed as follows:

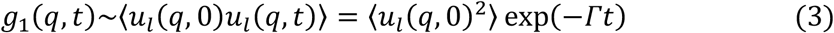

From *Γ* = *D*_gel_ *q*^2^ we can obtain the gel diffusion coefficient *D*_gel_, which is proportional to the longitudinal modulus *M* of the gel, *D*_gel_ = *M* /*f* (*f* is the friction coefficient between the gel strands and the solvent),^23, 24^ so that *D*_gel_ reflects the gel elasticity.

DLS data from PAm-PAc gels with 80% charge density, 0.5% crosslinking density (80c, 0.5xl), 1 M monomer concentration and 50 mM CaCl_2_ are presented as an example. The electric field correlation function *g*_1_(*q, t*) of the gel at scattering angle 30° is shown in Fig. 1a. It can be fitted by an exponential decay and a stretched exponential decay (red curve), and the quality of the fitting can be seen from the fitting residuals (blue triangles) in Fig. 1a. The corresponding relaxation time distribution function *f* (t) at scattering angle 30° can be obtained from CONTIN fitting method (Fig. 1b). The fast mode accounts for 77% of the total proportion. In Fig. 1c, the relaxation rates *Γ* obtained at different scattering angles are proportional to *q*^2^, and from the slope of *Γ*-*q*^2^ the gel diffusion coefficient of the fast mode *D*_1_ = (1.08±0.03)×10^−6^ cm^2^/s can be obtained.

**Fig. 1.**
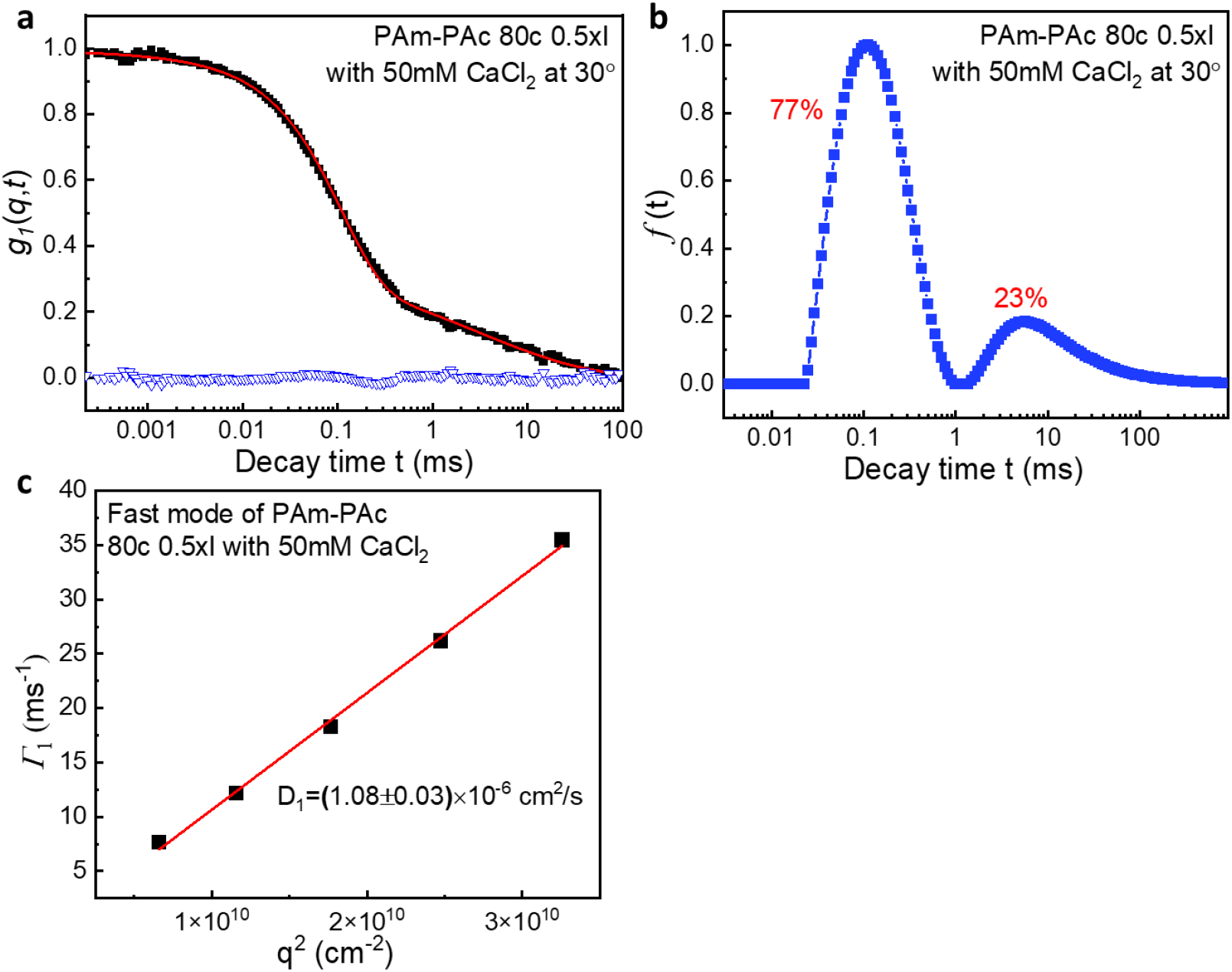
Dynamics light scattering measurement of PAm-PAc gel with 80c 0.5xl and 50 mM CaCl_2_. (a) Normalized electric-field correlation function *g*_1_(*q, t*) at scattering angle 30°. The red curve is the fitting curve and the blue open triangles are the residuals between the fitting curve and the raw data (black squares). (b) Relaxation time distribution function obtained from CONTIN fit at scattering angle 30°. (c) Corresponding *q*^2^ dependence of the relaxation rate *Γ* of the fast mode.

Concentration dependence of diffusion coefficient *D* was studied seen in Fig. 2. For 80c 0.5xl PAm-PAc gel without CaCl_2_, the correlation function has a diffusive mode. This mode indicates the elasticity of PAm-PAc chemical gel network, with *D*_0_ = (8.63±0.15)×10^−7^ cm^2^/s and 100% fraction (represented by green star) (Fig. 2a, b). When the gel is prepared containing even a very small amount of CaCl_2_, a slow mode emerges. For c(CaCl_2_)=5 mM, the fast mode accounts for 88.6±0.05% of the total fraction with *D*_1_ = (1.44±0.05)×10^−6^ cm^2^/s. The fast mode denotes the gel mode because of the similar order of magnitude with *D*_0_, so that the fast mode represents the elasticity of the chemical gel network. The change of c(CaCl_2_) has no influence on the volume of the as-prepared gels as the volume ratios of gels to pre-gel solutions are all around 0.97 seen in Fig. 3 and all the gels we measured are transparent at 23 °C. The gel will undergo phase separation at high salt concentrations represented by the shadow in Fig. 2. Unless otherwise specified, the changes in volume and appearance of the gels studied in this work share similar circumstances with the gels containing CaCl_2_. The transparency refers to that there are no micron phase domains in the gels. Besides, the gelation process occurs uniformly in space and rapidly in time to avoid any volume changes caused by osmotic pressure. When c(CaCl_2_) is below 100 mM, the chemical gel mode relaxes faster than that of c(CaCl_2_)=0, shown in Fig. 2a where *D*_1_ is larger than *D*_0_. As c(CaCl_2_) increases from 5 mM to 200 mM, *D*_1_ gradually decreases to one-third ((4.44±0.05)×10^−8^ cm^2^/s) and the value of *D*_1_ at c(CaCl_2_)=200 mM is half of *D*_0_. Moreover, we denote the slow mode as the dissociation-association hierarchical relaxations of the reversible physical crosslinks caused by the multivalent salts.^18^ The decrease of the fraction of fast mode from 100% at c(CaCl_2_)=0 to 35.6±3.5% at c(CaCl_2_)=200 mM indicates that the physical network expands along with the increase of c(CaCl_2_), and the relaxation of the physical crosslinks becomes the main mode at this concentration. It is expected that when c(CaCl_2_) is infinitely close to the phase separation concentration, the physical network may envelope most of the chemical network to generate macroscopic phase regions.

**Fig. 2.**
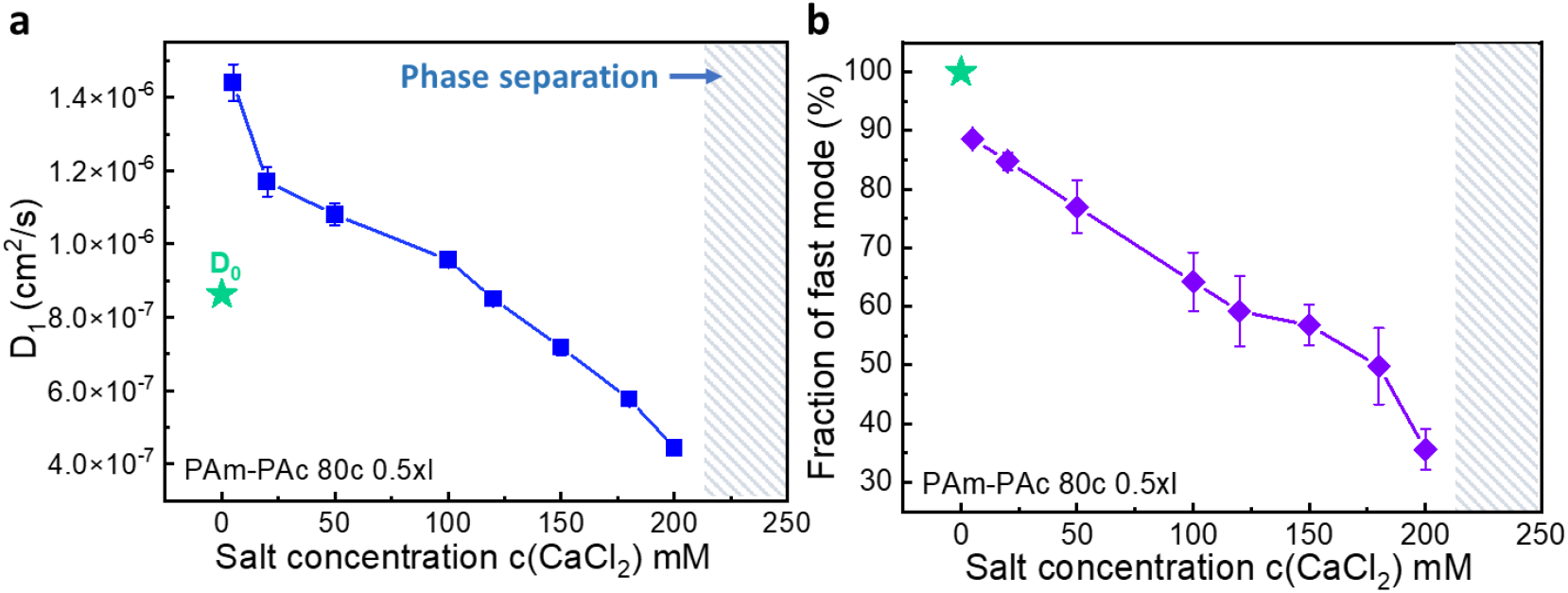
Dynamic light scattering results for PAm-PAc gels with 80c 0.5xl and different c(CaCl_2_). (a) Diffusion coefficients *D*_1_ of PAm-PAc gels with different c(CaCl_2_). (b) Fraction of the fast mode obtained by CONTIN fit.

**Fig. 3.**
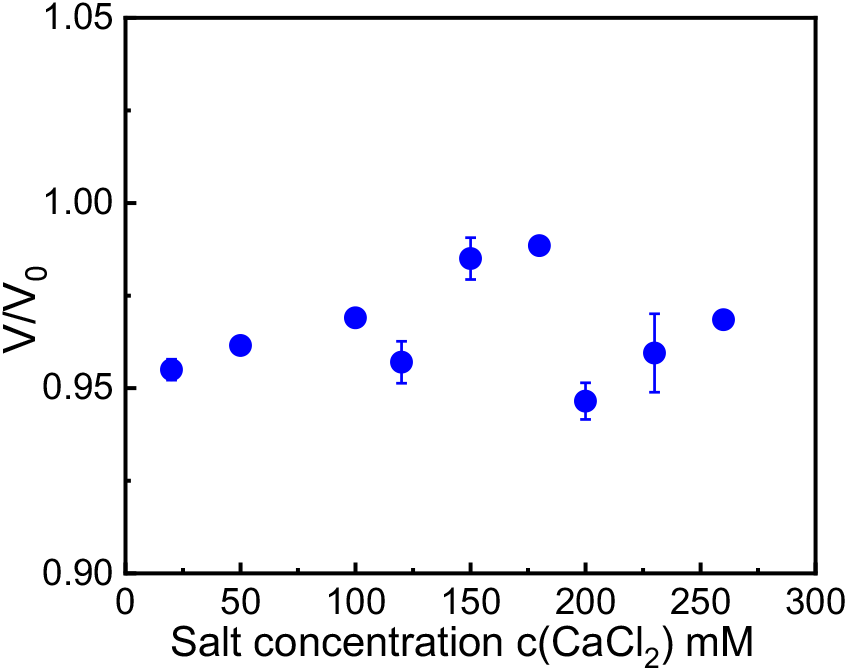
The ratio of the volume of PAm-PAc gels with 80c 0.5xl and different c(CaCl_2_) to the volume of the pre-gel solutions.

^1^H-NMR measurements were conducted to study whether calcium ions induce the decrease of the diffusion coefficient by impacting chemical conversion. In Fig. 4, peak a and b indicate the vinylic protons of Am monomers and Ac monomers respectively. 0.1 mol/L DMSO serves as an external standard substitute to determine the relative content of monomers in pre-gel solution and gels so that the chemical conversion can be obtained. Results show that the PAm-PAc gels with 50 mM, 150 mM and 250 mM CaCl_2_ have no difference with pure PAm-PAc gels in chemical conversion, which is around 96% for all the gels. It refers to the sufficient conversion of monomers. Therefore, CaCl_2_ should not decrease the diffusion coefficient of PAm-PAc gels through changing the chemical conversion.

**Fig. 4.**
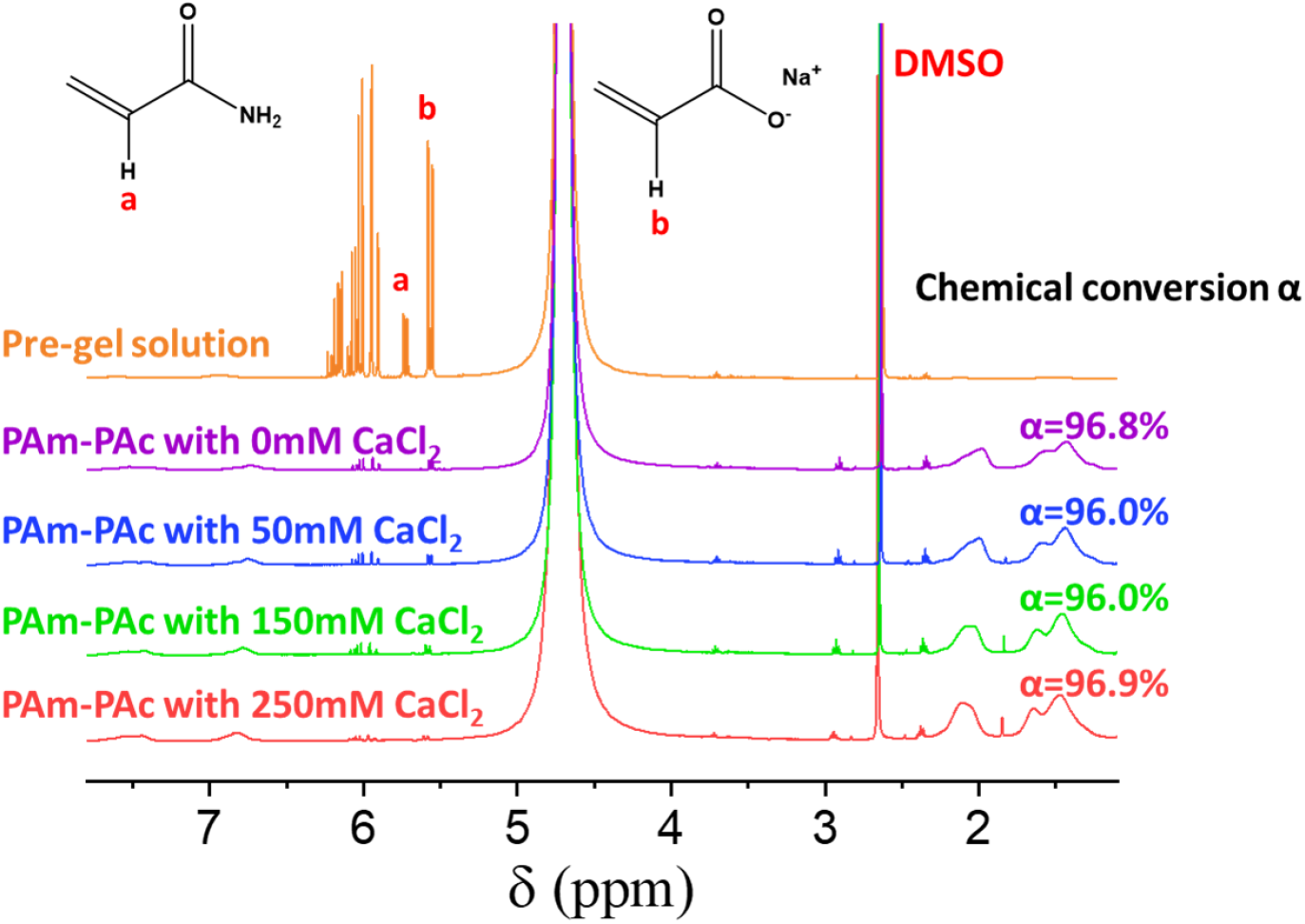
^1^H-NMR spectra of the pre-gel solution and PAm-PAc gels with 80c 0.5xl and different c(CaCl_2_).

In order to further understand these two modes of PAm-PAc gels, crosslinking density was changed from 0.2% to 1.0% as seen in Fig. 5. The as-prepared CaCl_2_ concentration should not be too high because the samples would not completely gel, for example, at c(CaCl_2_)=100 mM and 0.2xl. Moreover, the concentration should not be too low because the fractions of the fast mode among gels with different crosslinking densities would be very close to each other so that it would be difficult to identify the trend of fraction. For the PAm-PAc gel with 50 mM CaCl_2_, *D*_1_ fluctuates around 1.3×10^−6^ cm^2^/s from 0.2% to 1.0% (Fig. 5a). In Fig. 5c, a transition of the fraction can be seen between 0.35% and 0.5%, and from 0.65% to 1.0% the fractions of the fast mode maintain at around 80%. The increase of the fraction of fast mode along with the increase of the crosslinking density verifies that the fast mode represents the elasticity of the chemical gel network. Changing the crosslinking density cannot induce the diffusion coefficient to undergo a huge transition like the reduction of one order of magnitude induced by changing the concentration of CaCl_2_. Only the fraction of the fast mode is sensitive to the crosslinking density below a specific value. When the crosslinking density is high enough, we conjecture that the chemical network will dominate the gel structure so that the fraction of the fast mode remains unchanged.

**Fig. 5.**
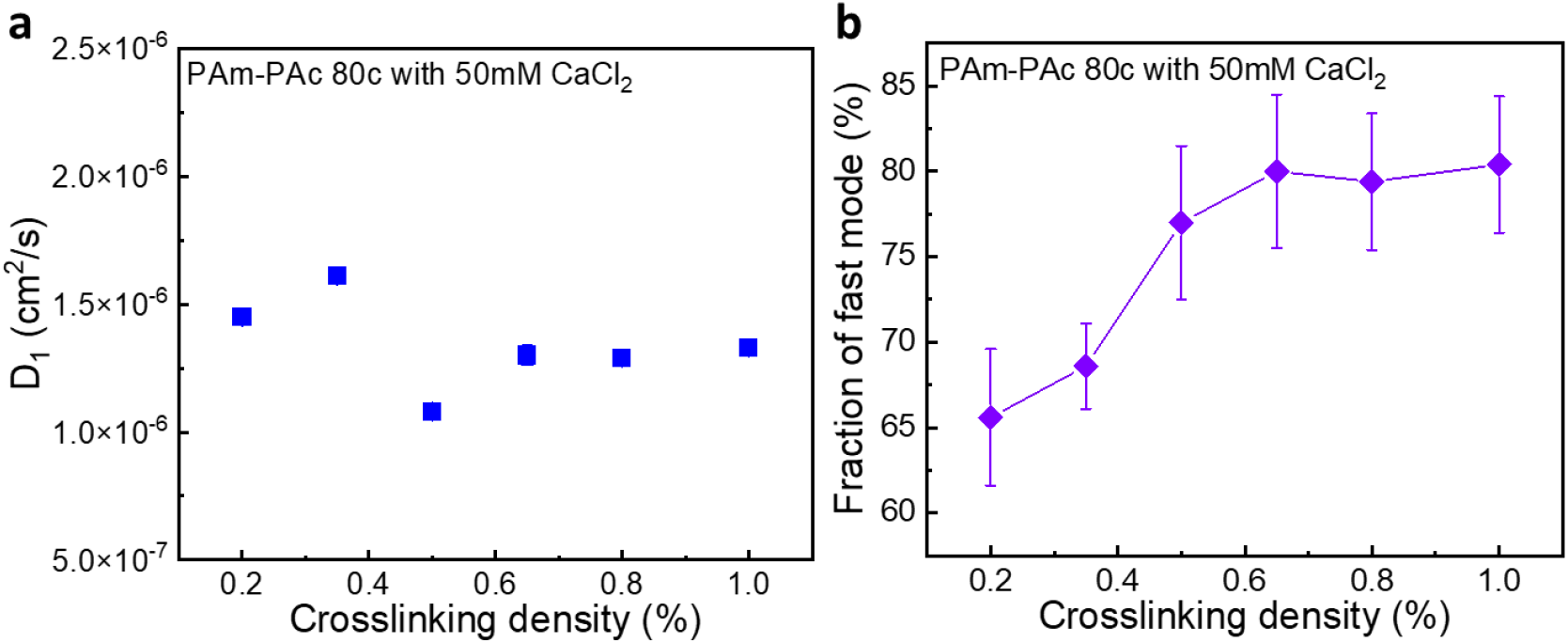
Dynamic light scattering results for PAm-PAc gels with different crosslinking density. (a) Diffusion coefficients *D*_1_ of PAm-PAc gels with different crosslinking density. (b) Fraction of the fast mode obtained by CONTIN fit.

To further study the multivalent salt induced transition in PAm-PAc gels, we introduced other types of cations and monitored the variation of the diffusion coefficient with concentration. Spermine is a kind of polyamine with relation to many biological activities such as cell proliferation. It has four positive charges at physiologic pH that binds strongly to acidic or negatively charged molecules in human body such as nucleic acids, or phospholipids ^7^. Before addition, the pH of spermine stock solution was adjusted to 6.86 to make it fully charged. Fig. 6a shows the chemical structure of spermine^4+^. The pre-gel solutions and as-prepared gels are all transparent within the spermine concentration range studied (0 – 250 mM), and volumes have no difference among gels with different c(spermine^4+^). The green stars refer to the original state of PAm-PAc gel without spermine^4+^. After addition of spermine^4+^, *D*_1_ and the fraction of fast mode decrease dramatically, and also a slow mode appears. For c(spermine^4+^)= 20 mM, *D*_1_ is almost the same value ((1.46±0.04)×10^−6^ cm^2^/s) as that of gel with c(CaCl_2_) = 5 mM ((1.44±0.05)×10^−6^ cm^2^/s) at 0.5xl. However, the fraction of fast mode at c(spermine^4+^)= 20 mM (58.8±4.5%) are much lower compared to c(CaCl_2_) = 5 mM (88.6±0.5%), which are close to c(CaCl_2_) = 120 mM (59.2±6%). To achieve the similar fraction of fast mode, the positive charges provided by Ca^2+^ at 120 mM are 3-fold higher than those of spermine^4+^ at 20 mM. It emphasizes that the important roles of chemical structure in the formation of physical network. When c(spermine^4+^) decreases to 100 mM, the fraction of fast mode reaches to its minimum of 31%. At this concentration, *D*_1_ and the fraction of fast mode both decrease by about three times. This trend has also been observed in gels with as-prepared Ca^2+^ (Fig. 2). Above 100 mM, *D*_1_ gradually approach to constant values, but the fraction of fast mode increases markedly to 66% at c(spermine^4+^)= 250 mM. We suppose that the carboxyl groups of Ac are charge saturated and the gels undergo charge reversal above 100 mM. Therefore, the fraction of fast mode increases with increasing c(spermine^4+^). Besides, we measured zeta potential to verify charge reversal induced by spermine^4+^. To ensure measurability of the studied systems, we polymerized PAm-PAc without crosslinker at different spermine^4+^ concentrations, and spermine^4+^ crosslinked PAm-PAc to form colloids making them viscous fluid. Samples are transparent at c(spermine^4+^) no more than 100 mM and undergo transition of zeta potential from negative to positive values, with the critical concentration being around 50 mM. When the c(spermine^4+^) is high enough, phase separation occurs and the zeta potential reverts to negative value. The deviation of critical point between gels and colloids may derive from the different local chemical environment of charged groups, which leads to different local ionized degrees.

**Fig. 6.**
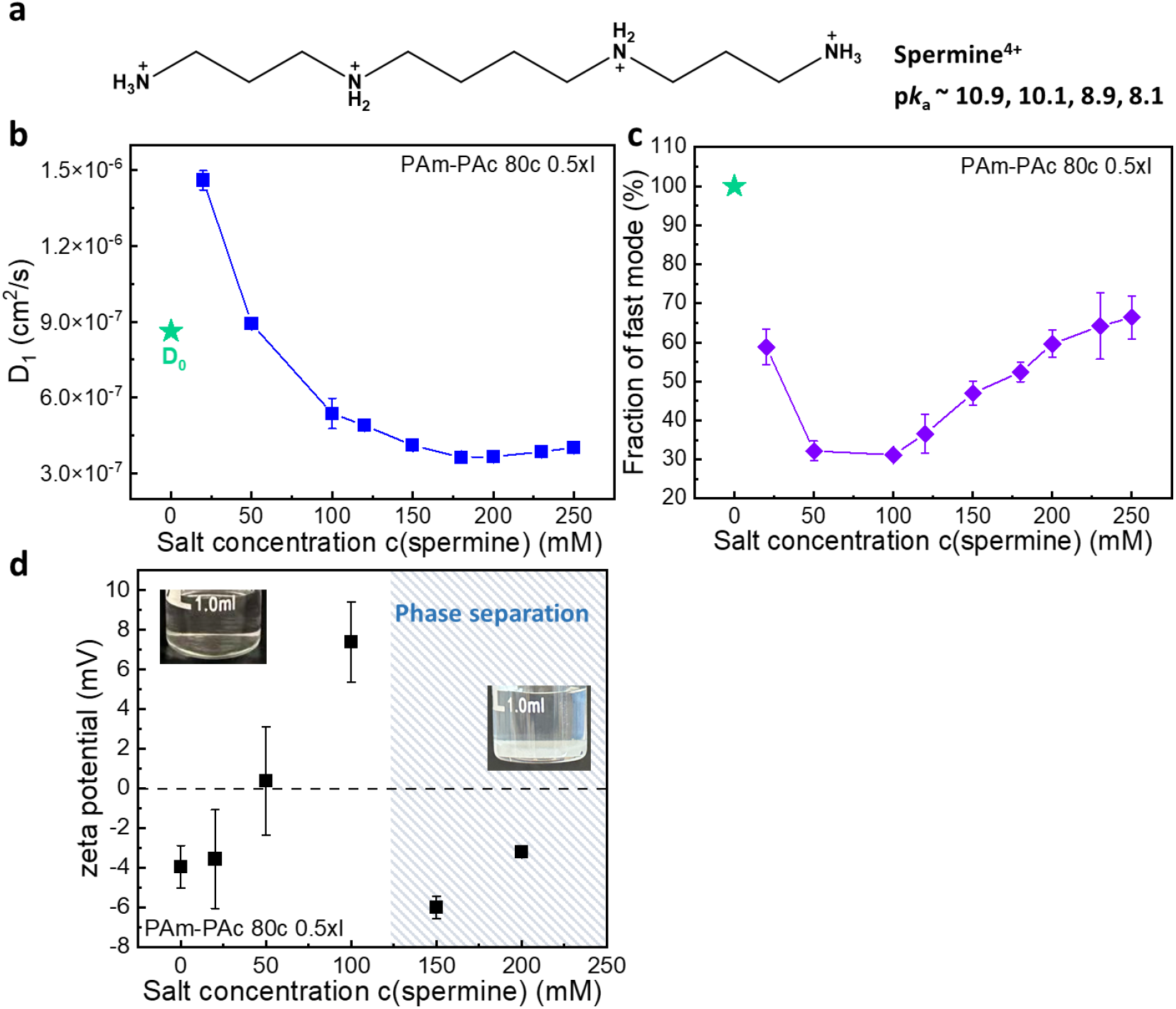
Concentration dependence of spermine^4+^. (a) Chemical structure of spermine^4+^. (b) Diffusion coefficients *D*_1_ of PAm-PAc gels with different c(spermine). (c) Fraction of the fast mode obtained by CONTIN fit. (d) c(spermine) dependence of zeta potential. PAm-PAc was polymerized without chemical crosslinker.

Multivalent counterions induced gel volume transition has been studied extensively. It is known that divalent cations with larger atomic mass induce anionic gel collapse at lower concentrations, and high-valent cations exhibit similar effect with lower transition points ^9, 25, 26^. However, in our research there seems to be no discernible mass dependence as *D*_1_ and the fraction of fast mode follow their own distinct patterns. In Fig. 7, several types of multivalent cations that would not hinder free radical polymerization are in situ prepared into PAm-PAc gels. Gels with low salt concentrations are featured with higher diffusion coefficients *D*_1_ that decrease with increasing salt concentration. At concentrations up to 20 mM greater than that of the final data point on each curve, the system undergoes phase separation, thereby rendering DLS measurement impossible. Among cations in Fig. 7, CaCl_2_ is least prone to cause phase separation while AlCl_3_ exhibits the strongest propensity that all the *D*_1_ measured are larger than *D*_0_. For each divalent cation, it can be seen that *D*_1_ around 100 mM is close to *D*_0_. It is worth noting that the spermine^4+^ concentration at which *D*_1_ gets close to *D*_0_ is 50 mM. So that for the alkaline earth metal cations studied and spermine^4+^, *D*_1_ will at first takes a large value, and then approach to its origin at the charge ratio [Ac^-^] /[M^n+^] = 4 (M^n+^ represents the cations mentioned above, and [Ac^-^] = 800 mM) as the concentration increases. As for the change of fraction, the slopes of MgCl_2_ and CaCl_2_ are similar while the BaCl_2_ dramatically decreases the fraction to 18.8% at 120 mM. From Fig. 7, it suggests that BaCl_2_ possesses a strong ability to aggregate Ac into physical networks and to further induced phase separation among alkali earth metal ions.

**Fig. 7.**
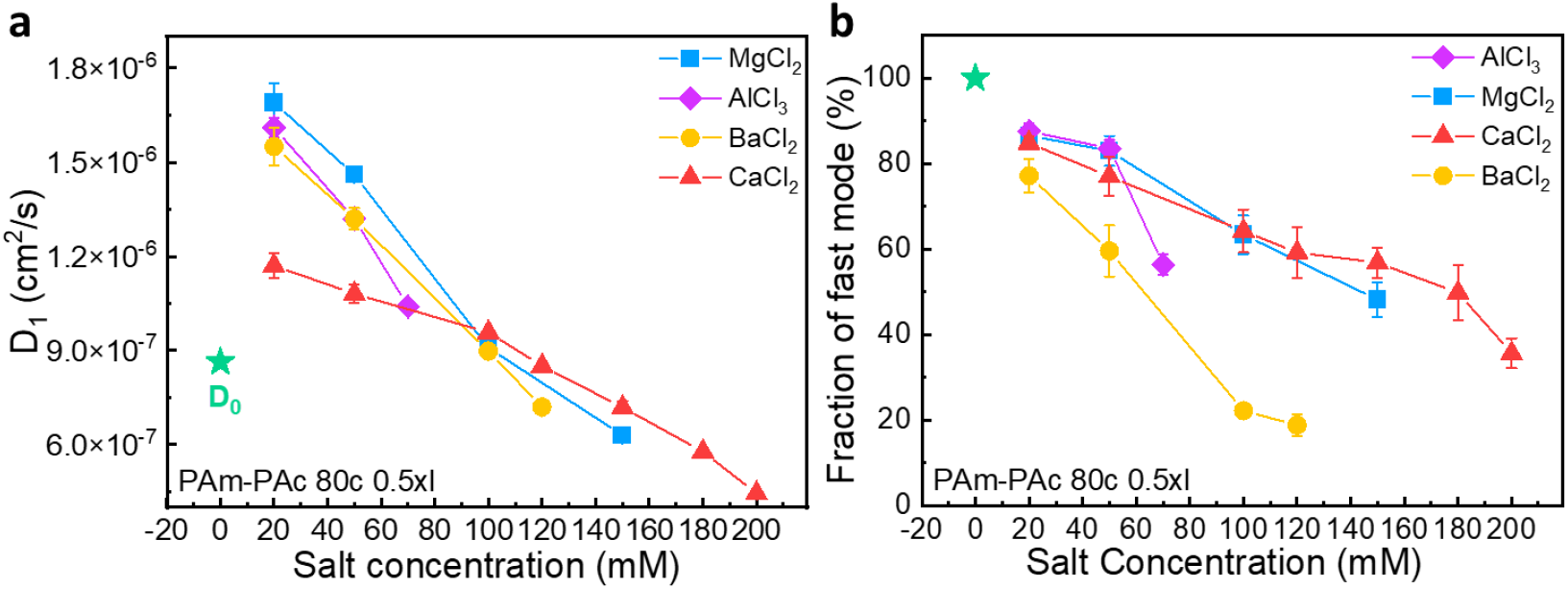
Effect of different multivalent salts. (a) Diffusion coefficients *D*_1_ of PAm-PAc gels with different salt concentration. (b) Fraction of the fast mode obtained by CONTIN fit.

As depicted in Scheme 1, PAm-PAc gel without multivalent salt is considered as a complete chemical gel, which is crosslinked by MBAA, a kind of organic small-molecule crosslinker (Scheme 1, left). When polymerization of PAm-PAc takes place in the presence of multivalent salts, the gel will be organized into two distinct regions so that it displays two relaxation modes in DLS. The fast mode corresponds to the chemical network because of the similar order of magnitude of diffusion coefficient. And we attribute the slow mode to the multivalent cations induced crosslinking of gel strands and the disassociation-association relaxion of the physical crosslinks (Scheme 1, right). In this case, the multivalent salts serve as crosslinkers making *D*_1_ larger than *D*_0_. Increasing multivalent salt concentration, the physical network expands in volume, which is dominant as the fraction is 70 ∼ 80% at high concentrations. The physical network significantly dissipates energy from thermal fluctuations through disassociation-association and any other deformation of strands in chemical network, combined with the condensation of ions onto the gel strands, leading to the decrease of *D*_1_.

**Scheme 1.**
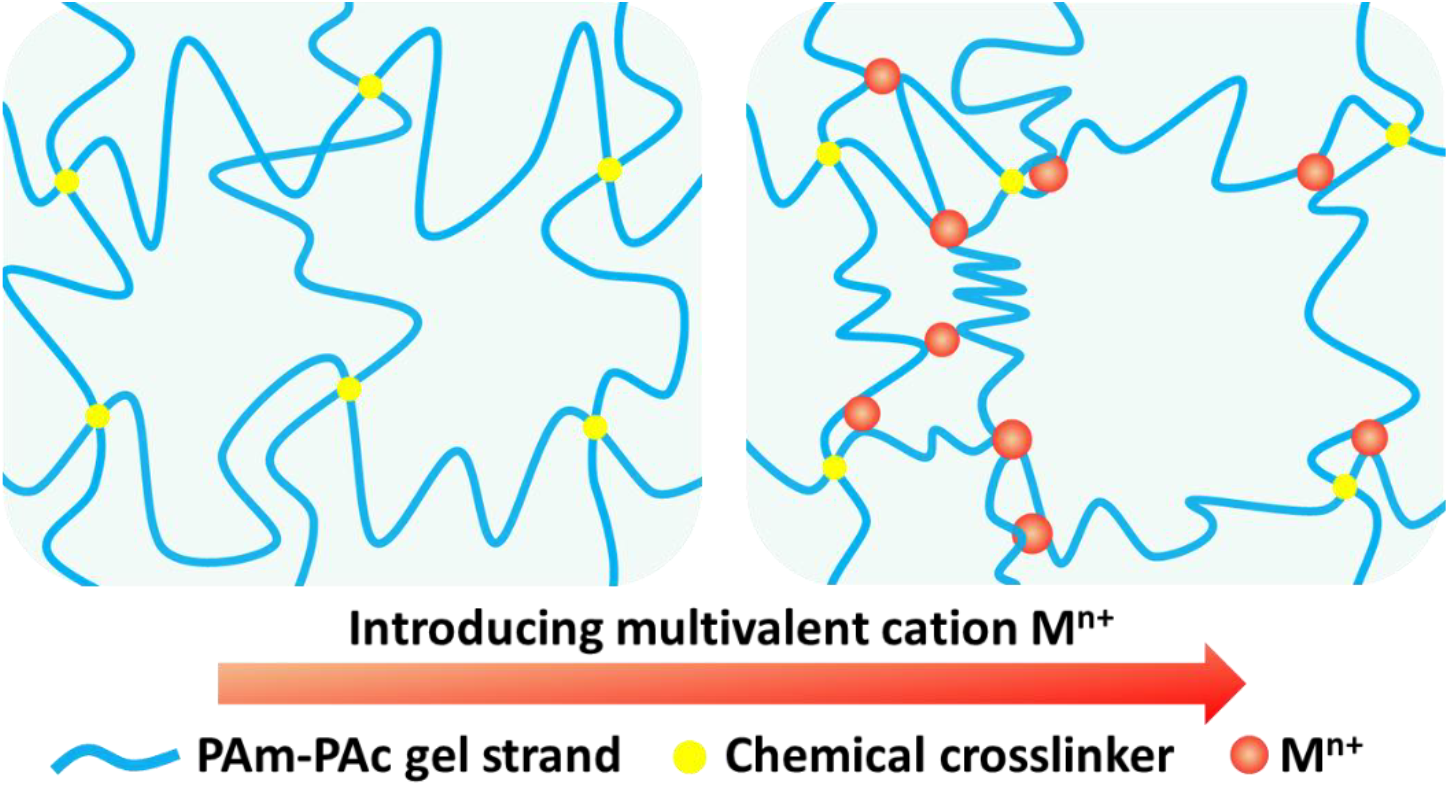
Schematic illustration of physical network induced by multivalent salt in chemical crosslinked gel.

## 4 Conclusions

In summary, we systematically investigate the multivalent cation effects on polyanion gels in their microscopic dynamics using dynamic light scattering. We prepare gels with series of multivalent cations featured by identical volumes and high chemical conversion. In this process, a new structure has been formed so that the gel is differentiated into two distinct yet interconnected parts: one is the chemical network and the other is physical network. They can be detected by DLS showing different relaxation modes, quantified by *D*_1_ and *D*_2_. For divalent and trivalent salts studied, increasing concentration will reduce *D*_1_, *D*_2_ and the fraction of fast mode monotonically. It indicates the slowdown of relaxation for two networks, the lower longitudinal modulus of gels, as well as the increasing weight of physical network. Moreover, *D*_1_ and *D*_2_ are hardly sensitive to chemical crosslinking density among a small range. In contrast, the fraction of fast mode shows sign of a growth plateau after a rapid increase with crosslinking density. In the case of high salt concentration, the gel is dominated by the physical network and primed for phase separation. For tetravalent salt spermine^4+^ with weak propensity to induce phase separation, we observe the charge reversal accompanied by a turning point of the fraction of fast mode. Our results provide basic understanding of multivalent ion-induced in-situ phase separation evolution in polyelectrolyte gels.

## ACKNOWLEDGEMENTS

This work was supported by the National Key R&D Program of China (Grant No. 2023YFE0124500), the National Natural Science Foundation of China (Grant No. 22273114), the Strategic Priority Research Program of the Chinese Academy of Sciences (Grant No. XDB0770101), and International Partnership Program of the Chinese Academy of Sciences (Grant No. 027GJHZ2022061FN).

